# The Impact of 2017 ACC/AHA Guidelines on the Prevalence of Hypertension and Eligibility for Anti-Hypertensive Therapy in the United States and China

**DOI:** 10.1101/218859

**Authors:** Rohan Khera, Yuan Lu, Anshul Saxena, Khurram Nasir, Harlan M. Krumholz

## Abstract

**BACKGROUND:** The 2017 American College of Cardiology (ACC)/American Heart Association (AHA) guideline recommendations for hypertension include major changes to the diagnosis of hypertension as well as suggested treatment targets for blood pressure management. To better guide future health policy interventions in the management of hypertension, we examined the effect of these guidelines on the prevalence as well as the eligibility for initiation and intensification of therapy in nationally-representative populations from the US and China.

**METHODS:** In the National Health and Nutrition Examination Survey (NHANES) for the most recent 2 cycles (2013-2014 and 2015-2016), and the China Health and Retirement Longitudinal Study (CHARLS) (2011-2012), we identified all adults 45 to 75 years of age who would have a diagnosis of hypertension, and would be candidates for initiation and intensification of anti-hypertensive therapy based on the 2017 ACC/AHA guidelines, compared with current guidelines.

**RESULTS:** The adoption of the 2017 ACC/AHA guidelines for hypertension in the US would label 70.1 million individuals in the 45-75-year age group with hypertension, representing 63% of the population in this age-group. The adoption of these guidelines in China would lead to labeling of 267 million or 55% individuals in the same age-group with hypertension. This would represent a relative increase in the prevalence of hypertension by 26.8% in the US and 45.1% in China with the adoption of the new guidelines. Further, based on observed treatment patterns and current guidelines, 8.1 million Americans with hypertension are currently untreated. However, this number is expected to increase to 15.6 million after the implementation of the 2017 ACC/AHA guidelines. In China, based on current treatment patterns, 74.5 million patients with hypertension are untreated, and is estimated to increase to 129.8 million if the 2017 ACC/AHA guidelines are adopted by China. In addition, the new ACC/AHA guidelines will label 8.7 million adults in the US, and 51 million in China with hypertension who would not require treatment with an anti-hypertensive agent, compared with 1.5 million and 23.4 million in the current guidelines. Finally, even among those treated with anti-hypertensive therapy, the proportion of undertreated individuals, i.e. those above target blood pressures despite receiving anti-hypertensive therapy and candidates for intensification of therapy, is estimated to increase by 13.9 million (from 24.0% to 54.4% of the treated patients) in the US, and 30 million (41.4% to 76.2% of patients on treatment) in China, if the 2017 ACC/AHA treatment targets are adopted into clinical practice in the respective countries.

**Conclusions:** Adopting the new 2017 ACC/AHA hypertension guidelines would be associated with a substantial increase in the prevalence of hypertension in both US and China accompanied with a marked increase in the recommendation to initiate and intensify treatment in several million patients. There would be a 26.8% and 45.1% increase in those labeled with hypertension in the US and China, respectively. Further, 7.5 million and 55.3 million will be newly recommended for therapy, and 13.9 million and 30 million newly recommended for intensification of existing therapy in the US and China, respectively.

## BACKGROUND

**T**he American College of Cardiology (ACC) and the American Heart Association (AHA) recently released guideline recommendations for treatment strategies for hypertension with lower blood pressure values used to define elevated blood pressure, and lower treatment thresholds,^1^ than those recommended in current guidelines.^2^ These guidelines define hypertension as a systolic blood pressure of 130 or greater or a diastolic blood pressure of 80 or greater, in contrast to a systolic blood pressure of 140 mmHg or greater or a diastolic blood pressure of 90 mmHg or greater that was used to define hypertension in all prior guidelines. Further, treatment recommendations have been extensively revised. The current guidelines recommend treatment of only a subset of patients with a diagnosis of hypertension. This includes all individuals with a systolic blood pressure of ≥140 mmHg or a diastolic blood pressure of ≥90 mmHg. Treatment was recommended for additional patients with lower blood pressures. Among those with a systolic blood pressure of 130-139 mmHg or a diastolic blood pressure 80-89 mmHg, treatment is recommended only for those who meet any of the following criteria: aged 65 years or older, pre-existing atherosclerotic cardiovascular disease (ASCVD) or a 10-year predicated risk of developing ASCVD of 10% or greater, chronic kidney disease (CKD), or diabetes mellitus. Further, a goal systolic blood pressure below 130 mmHg and a diastolic blood pressure below 80 mmHg is recommended for all patients, regardless of their baseline blood pressures. These recommendations contrast with systolic blood pressure targets of less than 140 mmHg and diastolic targets of less than 90 mmHg for most patients, other than targets of <150/90 mmHg for those ≥60 years of age without concomitant diabetes or CKD, suggested currently.^2^

The population impact of changes in guideline recommendations in the US are frequently evaluated. However, an understanding of how these recommendations translate to non-US populations, who frequently implement them, can also provide valuable insights into their overall impact. Further, since several major clinical studies rely on non-US populations, such international assessments of US guidelines are essential to appreciate their effect on other countries. China, in particular, has a high prevalence of hypertension and the world’s largest population. Recent studies suggest that even with current guidelines, a large proportion of adults in China are classified as hypertensive and few people are achieve blood pressure targets. Therefore, the impact of their adoption of these standards is important to understand.

Using nationally representative data from the United States and China, two diverse health systems, we examined the changes in prevalence, and eligibility for anti-hypertensive therapy and treatment intensification, based on current and the revised 2017 ACC/AHA guidelines.

## METHODS

### Data Sources and Study Population

We used contemporary data from large, nationally representative datasets from the US and China, which have been designed to assess the health status of community-dwelling individuals. For the US, we used the National Health and Nutrition Examination Survey (NHANES) for the most recent 2 cycles - 2013-2014 and 2015-2016, and for China, we used the baseline survey of the China Health and Retirement Longitudinal Study (CHARLS) conducted in 2011-2012.^3,4^ The structure of these datasets and their use to identify hypertensive adults have been described previously. Briefly, the NHANES, is a nationally representative database comprised of cross-sectional national surveys with information on demographic characteristics, medical history, prescription drug use, and laboratory testing on a set of individuals. The data in NHANES can be weighted to obtain nationally representative estimates. The CHARLS is a longitudinal study of 17,500 individuals >45 years of age from 150 counties/districts and 450 villages in China with a detailed baseline clinical evaluation in 2011-2012 when data on demographic characteristics, medical history, prescription drug use, and laboratory testing was collected.^3,4^ Further, similar to the NHANES, the data in CHARLS is nationally representative for China, and can be weighted to obtain national estimates for China.

In both these datasets, we identified all adults 45 to 75 years of age with at least two blood pressure measurements, and information on prescription drug use.

### Study Variables

From each dataset, we obtained information on all blood pressure measurements as well as variables necessary to classify patients into risk groups for anti-hypertensive therapy. This included information on demographic characteristics (age, sex, and race), clinical history (known ASCVD [defined as history of myocardial infarction, stroke, angina or reported coronary heart disease], smoking, diabetes [defined as hemoglobin A1c >6.5% or self-reported history of diabetes], and use of anti-hypertensive therapy) as well as laboratory values for serum creatinine, and total and high-density lipoprotein (HDL) cholesterol.

### Statistical Analysis

First, we calculated the average systolic and diastolic blood pressures based on all available readings for each patient. Second, we assessed the prevalence of hypertension, defined as the number of individuals who would have a diagnosis of hypertension based on current guidelines for both study populations. We use the 2014 evidence-based guidelines for the management of high blood pressure in adults developed by the panel members appointed to the eighth Joint National Committee as the ‘JNC-8’ criteria used in the context of the present study. These guidelines defined hypertension as an average systolic blood pressure of 140 mmHg or higher and/or a diastolic blood pressure of 90 or higher and/or treatment with an antihypertensive agent. We then identified individuals with hypertension based on the 2017 ACC/AHA guidelines. This included all adults meeting a definition based on the prior JNC-8 guidelines, but additionally included all adults with a systolic blood pressure 130-139 mmHg or a diastolic blood pressure of 80-89 mmHg. Next, we assessed the adults qualifying for treatment with one or more anti-hypertensive agents based on the current guidelines and the 2017 ACC/AHA guidelines but not receiving treatment (‘non-treatment’). This included all patients with a systolic blood pressure of 140 mmHg or higher or a diastolic blood pressure of 90 mmHg or higher. In addition, those with a systolic blood pressure of 130-139 mmHg or diastolic blood pressure of 80-89 mmHg were also treatment candidates if they were in one or more of high-risk groups: age ≥ 65 years, a history of ASCVD (defined as history of myocardial infarction, stroke, angina or reported coronary heart disease), CKD (based on an estimated glomerular filtration rate of <60 ml/kg/h calculated using serum creatinine and the Modification of Diet in Renal Disease study equation), diabetes [defined as hemoglobin A1c >6.5% or self-reported history of diabetes) or an expected 10-year risk of ASCVD of 10% or greater (defined using the pooled cohort equations).^5^ Finally, we assessed the number of patients who would be considered ‘under-treated’, based on being above the treatment targets defined in the older JNC-8 and the recent guidelines. In JNC-8, treatment threshold for hypertension were set at <150/90 mmHg for adults ≥60 years of age without diabetes or CKD, and <140/90 mmHg for all others. In the ACC/AHA guidelines, treatment targets were set at 130/80 for all patients. We used survey analyses with patient-level weights to obtain national estimates for both the US and China.

All statistical tests were two-sided and the level of significance was set at an alpha of 0.05. The study was exempt from the purview of Yale’s Institutional Review Board since it uses de-identified.

## RESULTS

### Prevalence of Hypertension: United States

In the NHANES dataset, a nationally-representative data from the United States, we found 2,973 participants, representing 55.3 million nationally (49.7% of the population) who are labeled with the diagnosis of hypertension based on the JNC-8 guidelines. An additional 685 individuals, representing 14.8 million (13.3% of the population) would be classified as hypertensive with the use of blood pressure cutoffs suggested in the 2017 ACC/AHA guidelines. Therefore, 70.1 million adults between 45 to 75 years of age would be classified as hypertensive based on the 2017 ACC/AHA guidelines, representing a 26.8% relative increase. The proportion of the US adult population between 45 and 75 years of age classified as hypertensive would increase from 49.7% based on the JNC-8 guidelines to 63.0% based on the 2017 ACC/AHA guidelines. The adoption of the new guidelines would be associated with a higher prevalence of hypertension across subgroups of sex and age. (**Figure 1**).

**Figure 1:**
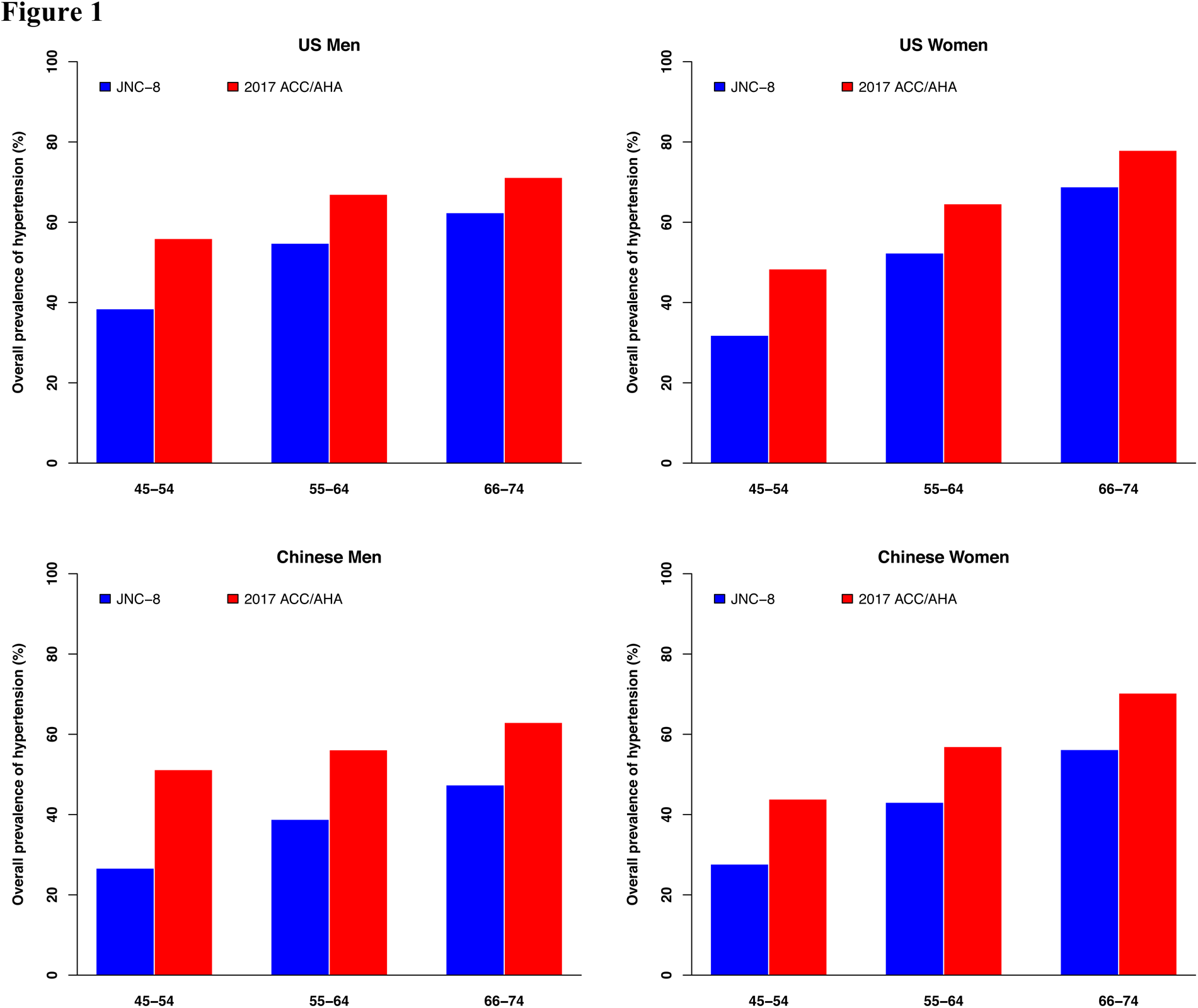
Prevalence of hypertension among study participants in the US and China, based on 2014 evidence-based guidelines for the management of high blood pressure in adults developed by the panel members appointed to the eighth Joint National Committee (JNC-8) and the 2017 ACC/AHA hypertension guidelines in the US (NHANES) and in China (CHARLS), among those between 45 to 75 years of age. Data are shown stratified by age and sex.

The characteristics of US patients labeled with hypertension according to the JNC-8 guidelines and the 2017 AHA/ACC guidelines are included in **Table 1**.

**Table 1:** Differences in characteristics of individuals with a diagnosis of hypertension based on 2014 evidence-based guidelines for the management of high blood pressure in adults developed by the panel members appointed to the eighth Joint National Committee (JNC-8) and the 2017 ACC/AHA hypertension guidelines in the US (NHANES) and in China (CHARLS), among those between 45 to 75 years of age.

| Characteristics, (%) | United States<br>(NHANES) |  |  | China<br>(CHARLS) |  |  |
| --- | --- | --- | --- | --- | --- | --- |
|  | JNC-8 | ACC/AHA | P-value | JNC-8 | ACC/AHA | P-value |
| **Number of people with hypertension** | 55,333,763 | 70,090,465 | <0.01 | 184,201,597 | 266,909,527 | <0.01 |
| **Demographics** |  |  |  |  |  |  |
| Age, y, median (IQR) | 61 (55-68) | 59 (52-66) | <0.01 | 59 (54-66) | 58 (52-65) | <0.01 |
| Sex |  |  | 0.92 |  |  | <0.01 |
| Men | 48.6% | 48.8% |  | 46.3% | 49.0% |  |
| Women | 51.3% | 51.2% |  | 53.7% | 51.0% |  |
| Race |  |  | <0.01 |  |  |  |
| Black | 14.1% | 13.2% |  | - | - | - |
| Non-black | 85.9% | 86.8% |  | - | - | - |
| **Comorbidities** |  |  |  |  |  |  |
| Diabetes | 26.7% | 23.1% | <0.01 | 17.4% | 15.5% | <0.01 |
| Smoking | 67.1% | 67.7% | 0.33 | 27.6% | 29.3% | 0.01 |
| CKD | 11.3% | 9.9% | <0.01 | 4.6% | 3.7% | <0.01 |
| ASCVD (MI, CHD, angina, stroke) | 15.9% | 13.4% | <0.01 | 19.4% | 16.1% | <0.01 |
| 10-year ASCVD risk, %, median (IQR) | 14.3 (8.3-23.2) | 12.6 (6.8-21.2) | <0.01 | 11.3 (5.6-20.4) | 9.6 (4.2-17.8) | <0.01 |
| **Labs, median (IQR)** |  |  |  |  |  |  |
| GFR | 85.6 (70.3-97.3) | 86.9 (72.2-98.1) | <0.01 | 92.5 (81.5-99.9) | 93.3 (82.6-100.9) | <0.01 |
| Total cholesterol, mg/dL | 192 (164-221) | 193 (168-224) | 0.08 | 193 (170-219) | 192 (170-217) | <0.01 |
| HDL cholesterol, mg/dL | 50 (41-63) | 51 (42-64) | 0.09 | 46 (38-56) | 47 (38-56) | <0.01 |
| Treatment with antihypertensive agents, % | 82.7% | 65.3% | <0.01 | 46.8% | 32.3% | <0.01 |

### Prevalence of Hypertension: China

In the CHARLS dataset, a nationally representative dataset from China, there were 5,133 individuals in the 45-75-year age-group who met the JNC-8 criteria for hypertension, representing 184 million (38.0% of the population) labelled with hypertension nationally. An additional 2588 individuals, representing 83 million (17% of population) nationally would be classified as hypertensive based on the 2017 ACC/AHA criteria. Therefore, 267 million Chinese adults between 45 and 75 years of age, would be classified as hypertensive based on ACC/AHA criteria, representing a relative increase of 45.1%. The proportion of the Chinese adult population between 45 and 75 years of age classified as hypertensive would increase from 38.0% based on the JNC-8 guidelines to 55% based on the 2017 ACC/AHA guidelines. Similar to the US, the adoption of the new guidelines would be associated with a higher prevalence of hypertension across subgroups of sex and age (**Figure 1**). The characteristics of patients with hypertension, based on the two criteria are included in **Table 1**.

### Eligibility for Anti-Hypertensive Therapy: United States

In the US, of the 55.3 million individuals with hypertension based on JNC-8 criteria, 82.7% were receiving one or more anti-hypertensive medications, while 14.6% (or 8.1 million individuals) were eligible for treatment but not receiving any therapy. Further, 1.5 million individuals in JNC-8 would be classified as hypertensive, and would qualify for lifestyle modifications alone. Of the 70.1 million classified as hypertensive based on the ACC/AHA guidelines, only 65.3% were receiving anti-hypertensive therapy, suggesting a recommendation to initiate therapy in 22.3% or 15.6 million individuals with hypertension (an additional 7.5 million individuals). Notably, under the 2017 ACC/AHA guidelines, the remaining 12.4% (or 8.7 million) of US adults in 45-75-year age-group would have hypertension, but not be candidates for anti-hypertensive therapy because they had a systolic blood pressure of 130-139 and/or a diastolic blood pressure 80-89 mmHg and without any of the high-risk conditions, such as age≥65 years, known ASCVD or a 10-year predicted risk of ASCVD of ≥10%, CKD, or diabetes. Moreover, of those receiving anti-hypertensive therapy, 24.0% were above the treatment targets for JNC-8 and 54.4% were above the treatment targets based on the ACC/AHA criteria. Therefore, an additional 13.9 million individuals will be candidates for intensification of therapy. The rates of non-treatment and under treatment of hypertension among subgroups of age and sex are presented in **Figures 2 and 3**.

**Figure 2:**
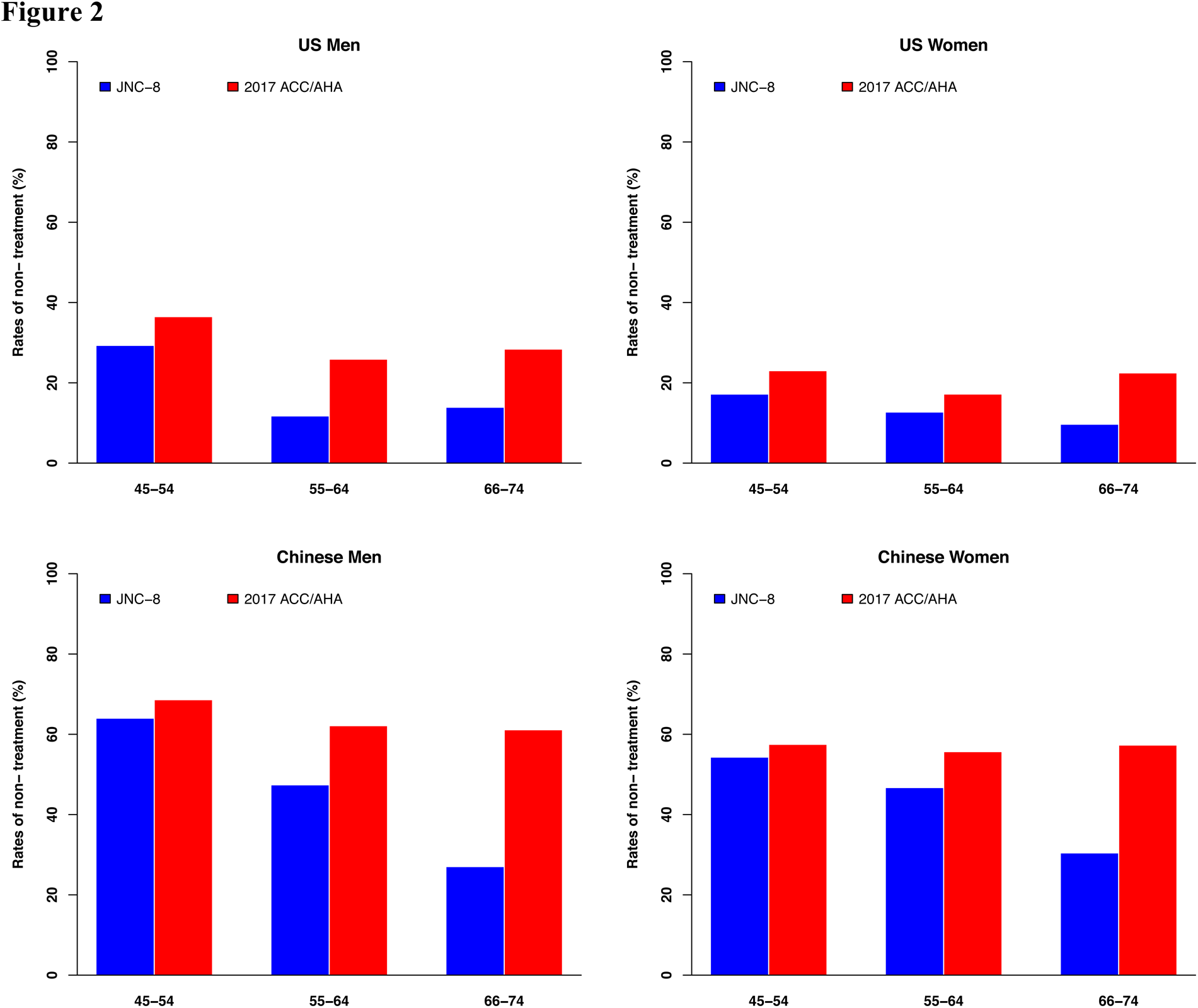
Rate of non-treatment among study participants with hypertension in the US and China, based on 2014 evidence-based guidelines for the management of high blood pressure in adults developed by the panel members appointed to the eighth Joint National Committee (JNC-8) and the 2017 ACC/AHA hypertension guidelines in the US (NHANES) and in China (CHARLS), among those between 45 to 75 years of age. Data are shown stratified by age and sex.

**Figure 3:**
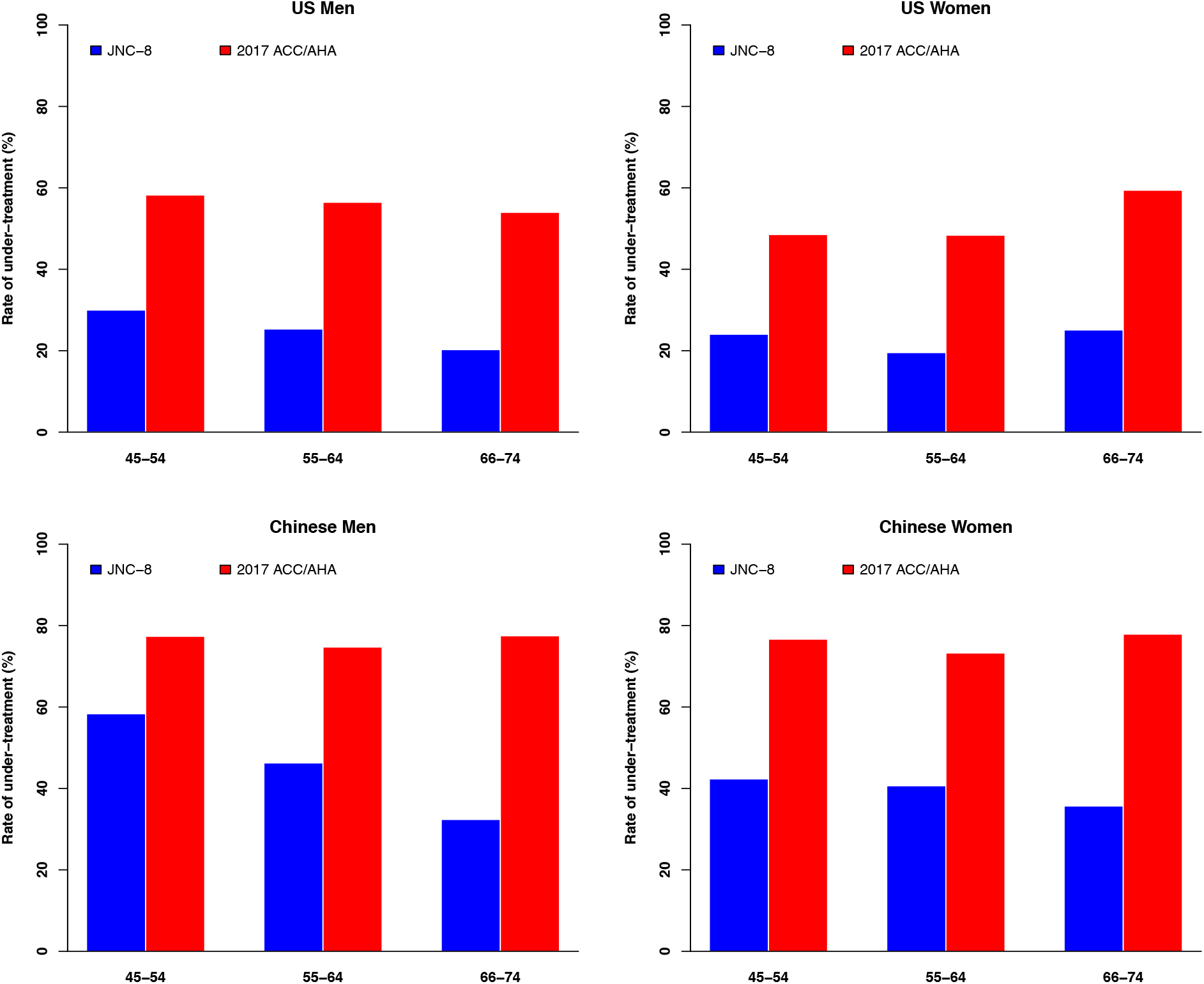
Rate of under-treatment among study participants with hypertension in the US and China, based on 2014 evidence-based guidelines for the management of high blood pressure in adults developed by the panel members appointed to the eighth Joint National Committee (JNC-8) and the 2017 ACC/AHA hypertension guidelines in the US (NHANES) and in China (CHARLS). Data are shown stratified by age and sex.

### Eligibility for Anti-Hypertensive Therapy: China

In China, based on JNC-8 blood pressure targets, 46.8% of the 184 million individuals with hypertension were receiving one or more anti-hypertensive medications. Among the 267 million classified as hypertensive based on the 2017 ACC/AHA guidelines, 32.3% were receiving anti-hypertensive therapy. The use of JNC-8 and 2017 ACC/AHA criteria to define treatment-eligible individuals would recommend the initiation of treatment in 74.5 million and 129.8 million patients (an additional 55.3 million individuals), respectively. Further, 23.4 million individuals in JNC-8 and 51.0 million individuals in the ACC/AHA guidelines would be classified as hypertensive, and would qualify for lifestyle modifications alone. This group included individuals with a systolic blood pressure of 130-139 and/or a diastolic blood pressure 80-89 mmHg and without any of the high-risk conditions, such as age≥65 years, known ASCVD or a 10-year predicted risk of ASCVD of ≥10%, CKD, or diabetes.

Finally, of those receiving anti-hypertensive therapy in China in 2011-2012, 41.4% were above the treatment targets for JNC-8, and 76.2% were above the treatment targets based on the ACC/AHA criteria. Therefore, an additional 30 million individuals would be candidates for intensification of therapy, based on the change in the guidelines. The rates of non-treatment and under treatment of hypertension among subgroups of age and sex are presented in Figures 2 and 3.

## DISCUSSION

The adoption of the 2017 ACC/AHA guidelines for hypertension in the US would classify 70.1 million individuals in the 45-75-year age group with hypertension, representing 63% of the population in this age group. The adoption of these guidelines in China would lead to the classification of 267 million or 55% individuals in the same group as suffering from hypertension. This would represent a relative increase of 26.8% in the US and 45.1% in China from prevalence based on current recommendations. Further, based on the observed treatment patterns and current guidelines, 8.1 million Americans with hypertension are currently untreated. However, this number is expected to increase to 15.6 million after the implementation of the 2017 ACC/AHA guidelines. In China, based on current treatment patterns, 74.5 million patients with hypertension are untreated and is likely to increase to 129.8 million if the 2017 ACC/AHA guidelines are adopted. Finally, even among those treated with anti-hypertensive therapy, the proportion of undertreated individuals, i.e. those above target blood pressures despite receiving anti-hypertensive therapy, is likely to increase by 13.9 million (from 24.0% to 54.4% of the treated patients) in the US, and 30 million (41.4% to 76.2% of patients on treatment) in China, if the 2017 ACC/AHA treatment targets are adopted into clinical practice.

The lowering of the treatment target for higher risk subgroups in the 2017 ACC/AHA is guided by evidence for a reduced rate of major adverse cardiovascular events.^6-8^ However, even based on the more conservative guidelines, the appropriate management of patients with anti-hypertensive therapy has posed major implementation challenges with high rates of non- and under-treatment in both the US and China.^9-11^ The ACC/AHA guidelines would require expansion of the public health infrastructure necessary to manage the substantial increase in the public health burden of hypertension in both the US and China. Further, health policy interventions focusing on appropriately treating recommended individuals would need to identify 7.5 million additional untreated Americans in the 45-75-year age-group, in addition to the 8.1 million who are not receiving treatment based on current recommendations. However, a similar effort in China would require identification of an additional 55.3 million untreated patients with hypertension, in addition to the 74.5 million currently with an unmet need for anti-hypertensive therapy. The rates of non-treatment, in combination with rates of under-treatment, suggest that health system interventions would need to address treatment in nearly all patients with hypertension.

Our study should be interpreted in the light of the following limitations. First, the data is based on cross-sectional surveys, with a potential for sampling error. However, since our estimates are based on a large subset of sampled individuals in both the NHANES and CHARLS, and are associated with a relative standard error of <10% for all estimates, the observations are likely to be reliable. Second, the assessment of national estimates was limited to adults 45 to 75 years of age, corresponding to the age group of individuals surveyed for the CHARLS study. Therefore, additional people <45 years and >75 years of age in both US and China may have hypertension, and be candidates for anti-hypertensive therapy. Third, while certain variables in both NHANES and CHARLS are based on self-reporting and are subject to recall bias. However, wherever appropriate we used objective data to define conditions (e.g. serum creatinine to assess CKD status) or complement the diagnoses derived from self-reporting (e.g use of A1c data to assess diabetes status). Further, data in both NHANES and CHARLS are collected by trained and experienced interviewers with the goal of reducing the effect of such biases. Fourth, a history of ASCVD in CHARLS includes those who may have congestive heart failure or other heart disease, in addition to prior myocardial infarction, stroke, coronary heart disease, and angina. Hence, the ASCVD rates are higher than the US. This did not affect classification with hypertension, which was solely based on an elevated blood pressure above 130/80 mmHg. A total of 3.7 million patients (i.e. 1.3% of overall patients) became treatment eligible due to ASCVD status in CHARLS. However, true ASCVD likely represented the vast majority of these patients, and the proportion of those with solely ‘congestive heart failure’ or ‘other heart disease’ is likely to be small. Fifth, we compared the recent guidelines against blood pressure definitions and blood pressure targets suggested by the members of the expert panel of the 8^th^ Joint National Commission, but physicians may follow different guidelines. However, hypertension has been defined consistently across all prior guidelines as a blood pressure ≥140/90 mmHg. Moreover, treatment targets in guidelines like the JNC-7 were more stringent,^12^ thereby suggesting an even higher rate of non-and under-treatment of hypertension in patients. Finally, we do not know if China will adopt these guidelines. Our study, however, only focuses on the potential impact of the uptake of these guidelines by China.

In conclusion, the adoption of the new 2017 ACC/AHA guidelines would be associated with a marked increase in the prevalence of hypertension in both US and China. While 6 million more US adults would require initiation of treatment, adoption in China would increase the number of treatment-eligible individuals by 31 million. There is a large unmet need for appropriate hypertensive therapy in China, and that challenge will grow further with the adoption of these guideline recommendations.

